# Involvement of monoaminergic and nitrergic pathways in the behavioral effects and acute toxicity of *Gomphrena perennis* L. tincture in mice

**DOI:** 10.64898/2025.12.26.696583

**Authors:** Adriana Milena Bonilla, Gabriela Garrido, Alicia E. Consolini, María Inés Ragone

## Abstract

*Gomphrena perennis* L. possesses a rich phenolic profile and has demonstrated cardiovascular and anti-inflammatory activities; however, its neuropharmacological properties remain unexplored. The behavioral effects, underlying mechanisms, and acute toxicity of *Gomphrena perennis* tincture (GphT) were investigated in mice. GphT showed no signs of acute toxicity. Furthermore, it did not alter the number of cross lines in the open field test (OFT), but it induced anxiogenic responses in the elevated plus-maze test (EPM) and the novelty-suppressed feeding test (NSFT), which were reversed by L-NAME, a non-selective nitric oxide synthases inhibitor. GphT also reduced immobility time in the forced swimming test (FST) and the tail suspension test (TST). This antidepressant-like effect was prevented by haloperidol (D1/D2 antagonist), ketanserin (5-HT2A/2C antagonist), L-NAME, and sildenafil (PDE5 inhibitor), while propranolol (β-blocker), prazosin (α1-an-tagonist), yohimbine (α2-antagonist), and ondansetron (5-HT3-antagonist) did not modify it. Therefore, GphT produced anxiogenic and antidepressant-like effects without impairing locomotion. These effects involve dopaminergic and serotonergic pathways and depend on nitric oxide-mediated signaling, suggesting that GphT exerts a modulatory influence on interconnected neurochemical systems.

## 1. Introduction

The genus *Gomphrena* (Amaranthaceae) contains approximately 120 species distributed throughout the America, Australia, and Indomalaya. Many species of this genus are widely used in traditional medicine to treat various disorders, including infectious, inflammatory, gastrointestinal and respiratory diseases, as well as liver and kidney disorders, urinary problems, and hypertension [1–4]. Scientific studies have demonstrated several of these traditional uses, reporting antimicrobial, antioxidant, cytotoxic, anti-inflammatory, analgesic, antiarthritic, antihyperalgesic, hypotensive, and antispasmodic effects in various *Gomphrena* species [5–8]. These findings scientifically support the traditional medicinal uses of the genus.

Particularly, *Gomphrena perennis* L. several properties have been documented, including emollient, diuretic, hemostatic, antidiarrheal, sedative and hypotensive effects [9–11]. Our previous preclinical studies support its traditional use as an antispasmodic and hypotensive plant mainly due to the inhibition of calcium influx pathways. Additionally, subacute oral administration of *Gomphrena perennis* tincture (GphT) induced cardioprotective effects against ischemia/reperfusion injury, a mechanism that involves NO production [6].

Chromatographic analysis of GphT by RP-HPLC-DAD revealed a complex phenolic composition, highlighting flavonoids, particularly glycosylated flavones [6]. Besides, we identified compounds including glycosylated derivatives of apigenin and luteolin, as well as the flavanones eriocitrin and neoeriocitrin. The presence of (+)-catechin, phenolic acids (gallic and p-coumaric), isovitexin, and diosmin was also confirmed [6]. Ethnobotanical and phytochemical reports of the genus have identified various bioactive compounds, such as flavonoids, saponins and ecdysteroids, particularly 20-hydroxyecdysone (20-HE) [12–14]. This compound is also present in related Amaranthaceae, including *Pfaffia glomerata* and *Achyranthes aspera*, both of which have shown anxiolytic and antidepressant effects in animal models [15, 16]. Taken together, these findings highlight the pharmacological potential of *Gomphrena perennis* and support current research, which focuses on the central effects of GphT and the involvement of monoaminergic and nitrergic pathways.

Therefore, this study aimed to assess the central effects of *Gomphrena perennis* L. tincture (GphT) in behavioral paradigms and to investigate the mechanisms involved. We also assessed its acute toxicity in mice via intraperitoneal administration.

## 2. Results

### 3.1. Acute toxicity study of *Gomphrena perennis* tincture

Mice were treated intraperitoneally with *Gomphrena perennis* tincture (GphT) at 5, 50, 300 and 2000 mg/kg. No deaths were recorded at any dose. No changes in body weight were observed (Figure 1), and no evident signs of toxicity were detected (Table 1). Therefore, the doses of GphT used in the in vivo behavioral studies were set at 400 mg/kg and 800 mg/kg for subsequent pharmacological evaluation. These doses have been previously used to evaluate the central nervous system action of extracts from phylogenetically related species [19, 20].

**Figure 1.**
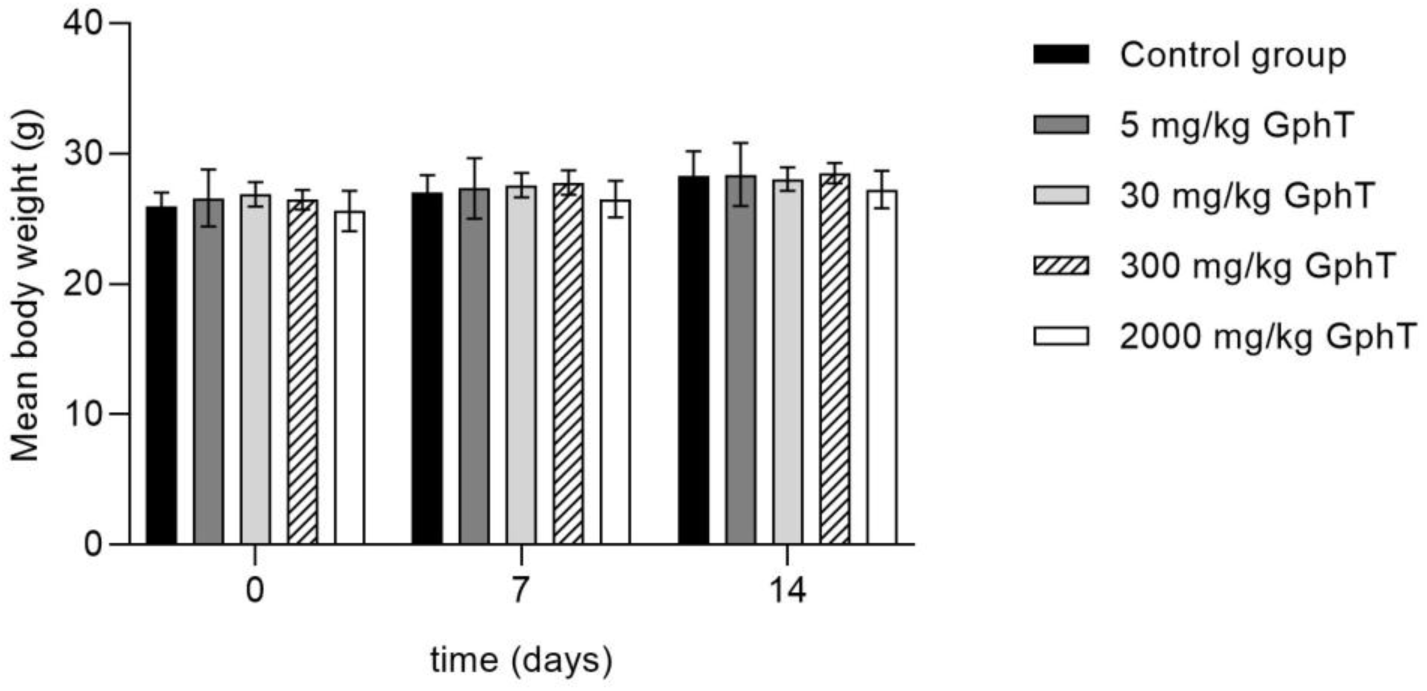
Body weight assessment of mice (n=3) treated with *Gomphrena perennis* tincture (GphT) during the acute toxicity study. One-way ANOVA: F = 0.7613, p > 0.5.

**Table 1.**
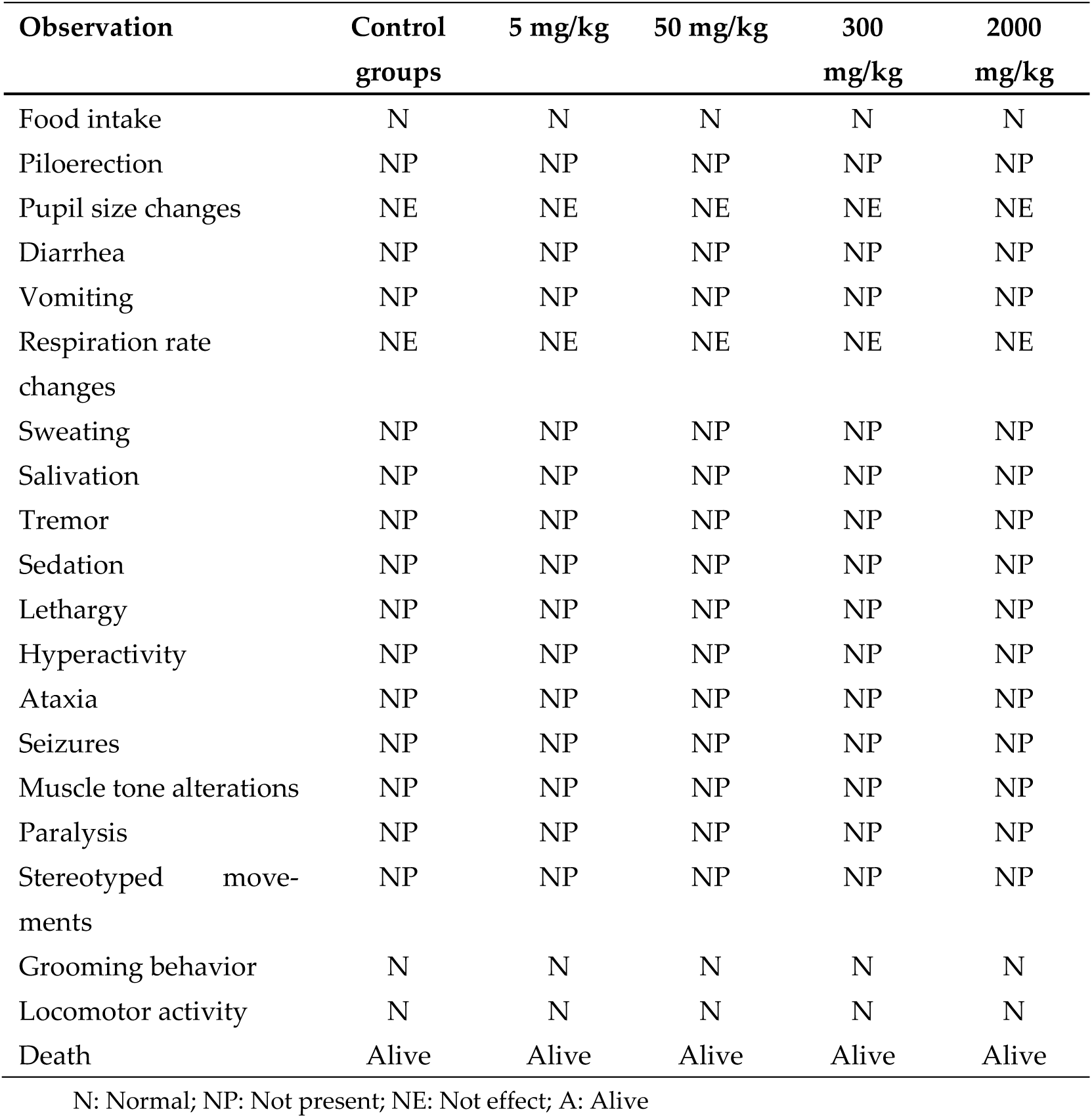
Table 1: Behavioral responses and general appearance of mice (n=3) after a single-dose administration of *Gomphrena perennis* tincture (GphT) in the acute toxicity study.

| Observation | Control groups | 5 mg/kg | 50 mg/kg | 300 mg/kg | 2000 mg/kg |
| --- | --- | --- | --- | --- | --- |
| Food intake | N | N | N | N | N |
| Piloerection | NP | NP | NP | NP | NP |
| Pupil size changes | NE | NE | NE | NE | NE |
| Diarrhea | NP | NP | NP | NP | NP |
| Vomiting | NP | NP | NP | NP | NP |
| Respiration rate changes | NE | NE | NE | NE | NE |
| Sweating | NP | NP | NP | NP | NP |
| Salivation | NP | NP | NP | NP | NP |
| Tremor | NP | NP | NP | NP | NP |
| Sedation | NP | NP | NP | NP | NP |
| Lethargy | NP | NP | NP | NP | NP |
| Hyperactivity | NP | NP | NP | NP | NP |
| Ataxia | NP | NP | NP | NP | NP |
| Seizures | NP | NP | NP | NP | NP |
| Muscle tone alterations | NP | NP | NP | NP | NP |
| Paralysis | NP | NP | NP | NP | NP |
| Stereotyped movements | NP | NP | NP | NP | NP |
| Grooming behavior | N | N | N | N | N |
| Locomotor activity | N | N | N | N | N |
| Death | Alive | Alive | Alive | Alive | Alive |

N: Normal; NP: Not present; NE: Not effect; A: Alive

### 3.2. Anxiolytic and sedative effects of *G. perennis* L. tincture on mice

GphT (400 and 800 mg/kg, i.p.) significantly reduced the time spent in the open arms of the elevated plus-maze test (EPM) compared to the vehicle (Figure 2a), without affecting the number of arm entries (Figure 2b). This effect was statistically significant only at the highest dose (800 mg/kg). In contrast, diazepam, a well-known benzodiazepine with anxiolytic properties, significantly increased both the time spent in the open arms and the number of arm entries compared to saline. Moreover, GphT at 800 mg/kg significantly increased the latency to eat in the novelty-suppressed feeding test (NSFT) compared to vehicle which was reversed by pretreatment with L-NAME (10 mg/kg, a non-selective nitric oxide synthase (NOS) inhibitor) (Figure 2c). Conversely, diazepam significantly reduced the latency to eat compared to saline. No change were recorded in home-cage food consumption for either treatment (Figure 2d).

**Figure 2.**
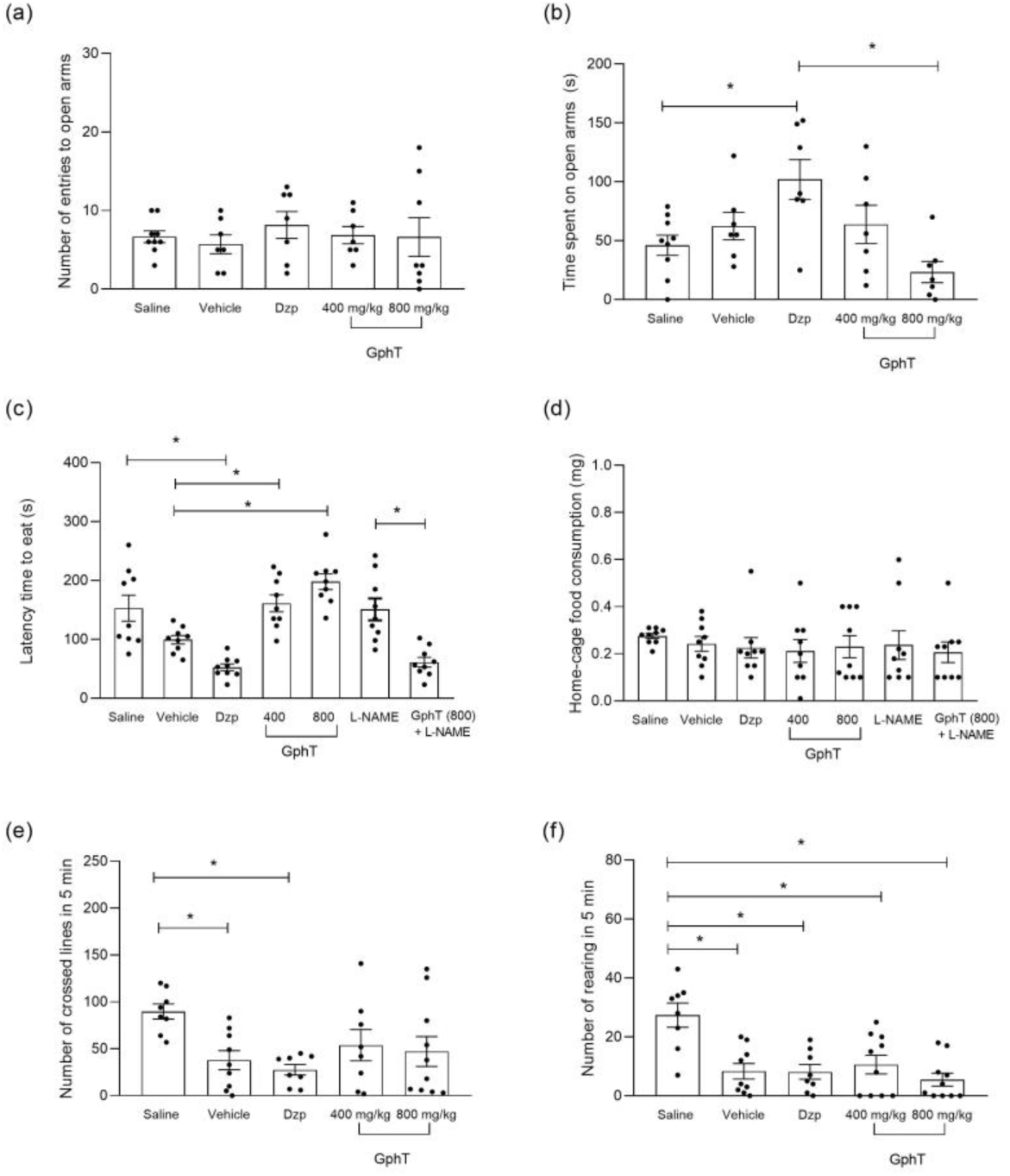
Behavioural tests evaluating the anxiolytic– and sedative-like effects of Gomphrena perennis tincture (GphT) in mice: time and entries in open arms. (a, b) in the elevated plus maze (EPM); latency to eat and home-cage food consumption (c, d) in the novelty-suppressed feeding test (NSFT); number of crossed lines and rearings (e, f) in the open field test (OFT). Data are expressed as mean ± SEM with individual values. One-way ANOVA followed by Tukey’s post hoc test: *p < 0.05 (Table A1).

To evaluate the possible sedative effect of GphT, the two doses were tested using the Open Field Test (OFT). GphT did not affect spontaneous locomotion, measured by the number of lines crossed in 5 minutes (LC), nor exploratory behavior, measured by the number of rearings in 5 minutes (RE), compared to the vehicle group. No significant differences were observed among the mice in the parameters measured in the OFT prior to treatment (Figure 2e and 2f).

### 3.4. Antidepressive effects of *G. perennis* L. tincture on mice and mechanism of action involved

The antidepressant effects of GphT were assessed using the TST and FST. Immobility time (IT) was measured in both experimental models, as a reduction in IT is widely accepted as a well-established measure of antidepressant-like activity, indicating a decrease in behavioral despair under conditions of inescapable stress [24]. IT was significantly reduced in mice treated with both doses of GphT (400 mg/kg and 800 mg/kg), similar to clomipramine in both the TST (Fig 3a) and the FST (Figure 3b). There were no changes in the number of crossed lines in the OFT in any treatment (Figure 3c)

**Figure 3.**
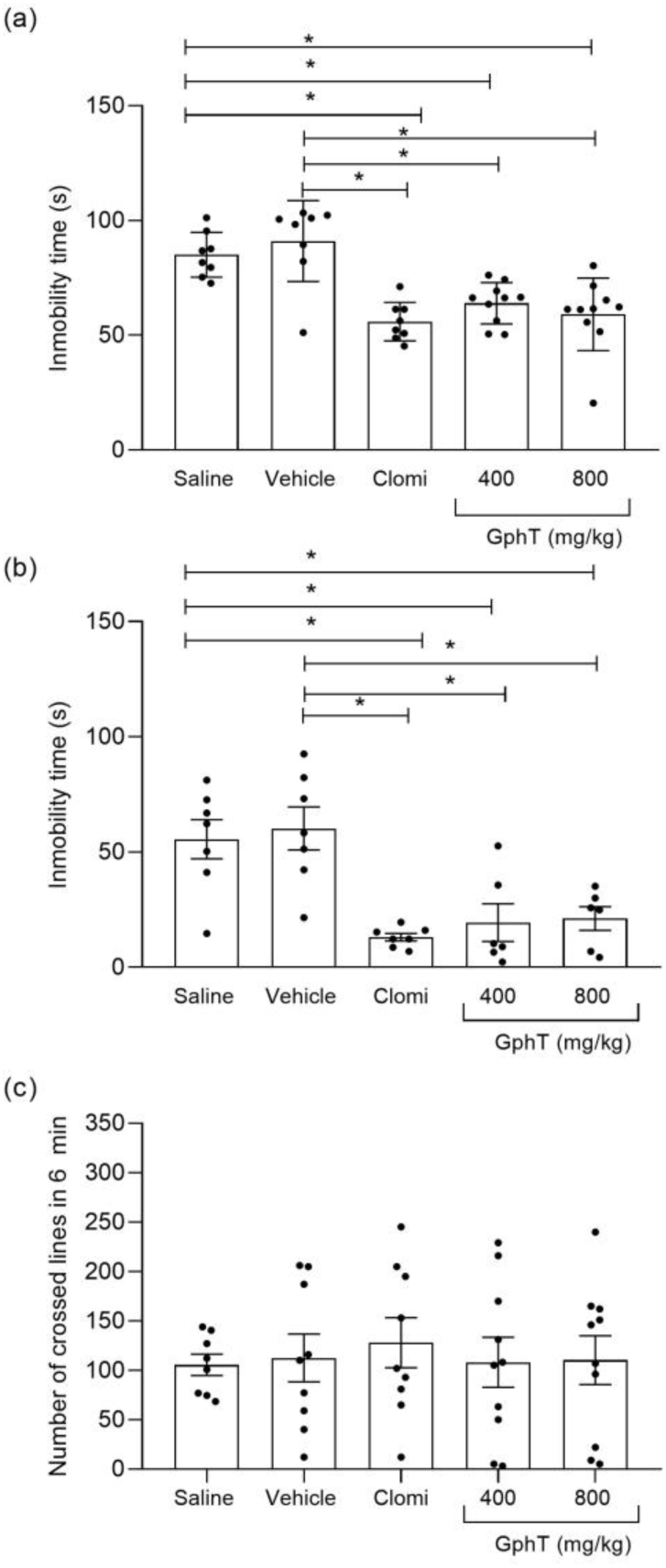
*Gomphrena perennis* tincture (GphT) effects on duration of immobility in Tail Suspension Test. (a) and in Force swimming test (b) and on number of crossed lines in 6 min in Open field test (c). Data are expressed as mean ± SEM with individual values. One-way ANOVA followed by Tuckeýs post hoc test: *p < 0.05 (Table A2).

Following pretreatment with monoaminergic receptor antagonists, ketanserin and haloperidol were able to reverse the antidepressant-like effect of GphT at 800 mg/kg, as evidenced by a significant increase in immobility time (IT) in the TST (Fig. 4d and 4e), Contrarily, neither propranolol (Fig. 4a), prazosin (Fig. 4b), yohimbine (Fig. 4c), nor ondansetron (Fig. 4f) modified it.

**Figure 4.**
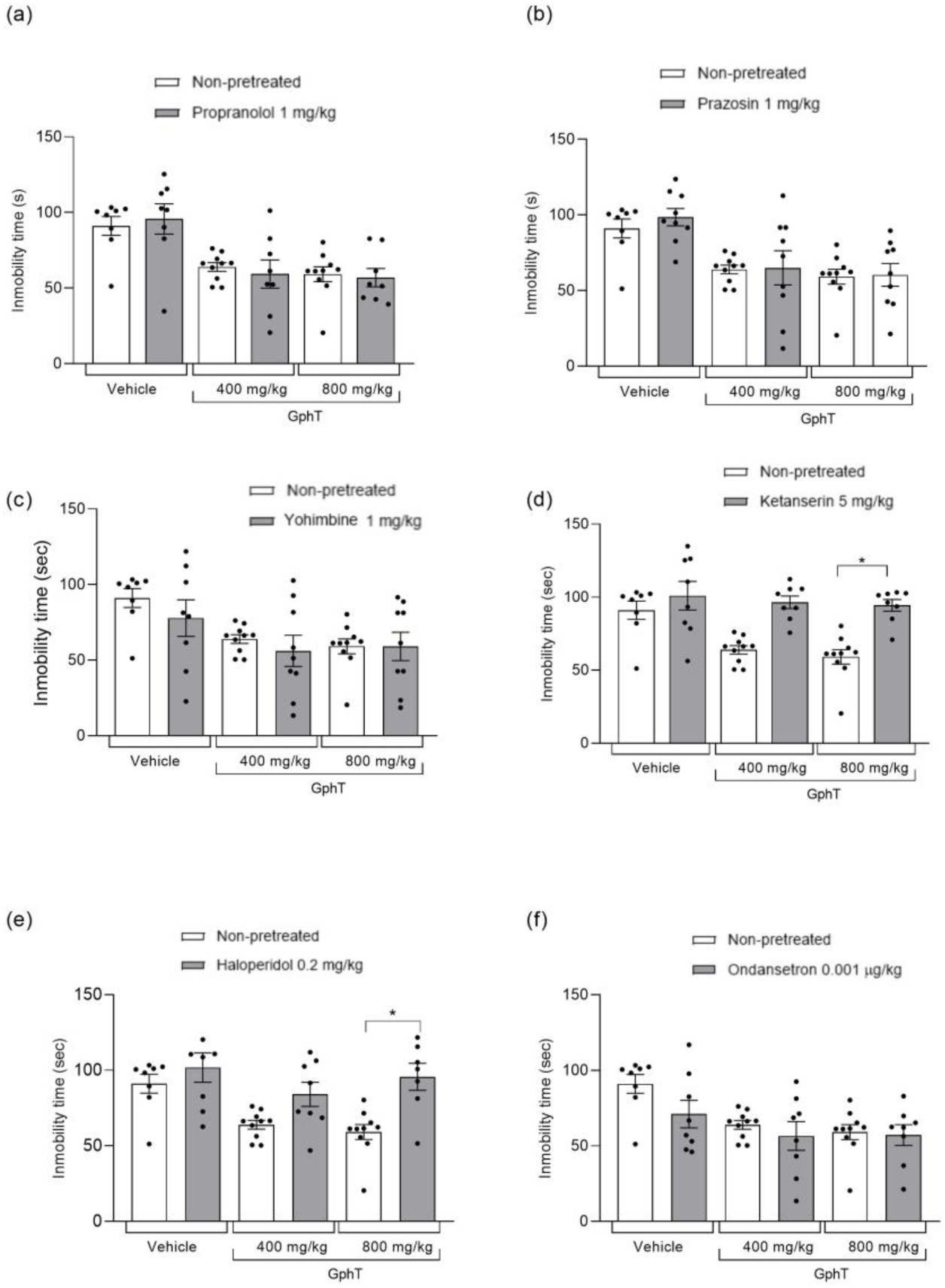
Effects of pretreatment with propranolol. (a), prazosin (b), yohimbine (c), ketanserin (d), haloperidol (e) and ondansetron (f) before Gomphrena perennis tincture (GphT) in Tail Suspension Test (TST). Data are expressed as mean ± SEM with individual values. Two-way ANOVA: P < 0.001 (Table A3). A posteriori Tukeýs test: *p < 0.05 vs. non-pretreated.

In addition, to evaluate the involvement of the nitric oxide (NO) pathway in the antidepressant-like effects of GphT, L-NAME was used in the forced swimming test (FST), where the role of NO in antidepressant-like responses has been consistently reported [31, 32, 33]. L-NAME reversed the effect of GphT by increasing IT (Figure 5a). In addition, the PDE5 inhibitor sildenafil, was used to investigate the role of the cGMP pathway in the antidepressant-like effects of GphT [31]. Sildenafil increased the immobility time of both doses of GphT, thus impairing its antidepressant effect (Figure 5b). There was no change in the number of crossed lines at 6 min in de OFT in mice treated with selective blockers in the presence of vehicle or GphT. (Fig A1, supplementary data).

**Figure 5.**
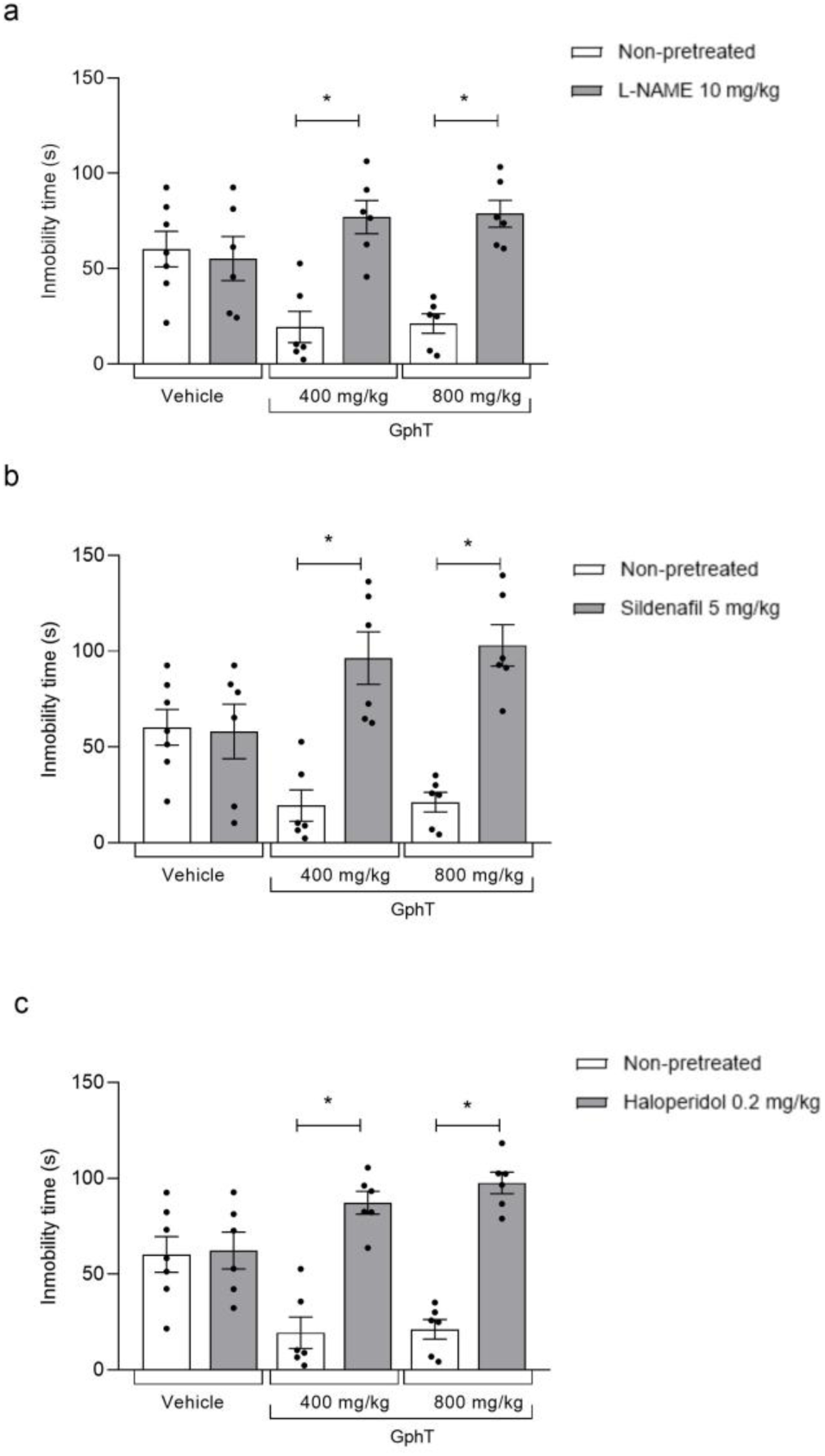
Effects of pretreatment with L-NAME. (a), sildenafil (b) and haloperidol (c) before Gomphrena perennis tincture (GphT) in Force swimming test (FST). One-way ANOVA: F = 0.7613, p > 0.5.

## 3. Discussion

This is the first study to provide scientific evidence of the central effects of *Gomphrena perennis* L. tincture (GphT) evaluated in preclinical behavioral models. Although *Gomphrena* species are widely used in traditional medicine for the treatment of various pathologies [6, 12], there are no previous studies evaluating their neuropharmacological profile. In the present work, we investigated the potential sedative, anxiolytic, and antidepressant-like actions of GphT, as well as its acute toxicity and the mechanisms of action involved. These findings are discussed in the context of existing literature and possible mechanisms of action.

Since no signs of acute toxicity were observed with intraperitoneal administration of 2000 mg/kg of GphT, we studied the behavioral effects of this extract through in vivo studies in mice. GphT did not exhibit sedative effects in the open field test, maintaining the number of lines crossed similar to vehicle. However, it induced anxiogenic response in both the elevated plus-maze and the novelty-suppressed feeding tests, evidenced by a low time spent on open arms and a significantly increased latency to eat, respectively. This anxiogenic effect was reversed by L-NAME, a non-selective inhibitor of nitric oxide synthase (NOS), suggesting that high levels of nitric oxide (NO) would be responsible at least in part for this effect.

Our finding contrast with bibliographic reports about anxiolytic and sedative effects of some of the components we have previously identified in GphT tincture by RP-HPLC-DAD (see supplemental data, Appendix A1) [6]. Among them, gallic acid has been described as anxiolytic at low doses and sedative at higher doses [34]. Similarly, (+)-catechin have been identified in extracts of sedative plants in preclinical studies [35–38]. Although baicalin has shown anxiolytic and sedative effects [39], other studies suggest a dissociation between its anxiolytic action and motor or sedative effects, possibly due to preferential affinity for GABA(A) receptor subtypes containing α2 and α3 subunits—related to anxiolysis—rather than α1 subunits, which mediate sedation [40, 41].

The lack of the expected pharmacological effect in GphT, despite containing known bioactive compounds, may be explained by complex interactions between among its components, differences in pharmacokinetics and bioavailability, variations in the molecular selectivity, heterogeneity of the plant material, and differences the experimental designs.

Furthermore, our findings show that nitric oxide (NO) is critically involved in the anxiogenic response. The involvement of NO in the subacute GphT treatment has been previously reported as part of the cardioprotective mechanisms [6]. It is widely recognized that the role of NO in the regulation of anxiety is bidirectional and highly dependent on the physiological context. Several studies have shown that inhibition of NOS generates anxiolytic responses in animal models [42, 43]. However, L-NAME has also been reported to induce anxiogenic responses in the elevated plus-maze test. This dual behavior supports the concept that NO acts as a bidirectional neuromodulator, whose effect—anxiolytic or anxiogenic—depends on multiple variables such as regional NOS expression, local NO concentration, the neuronal circuits involved, and the behavioral paradigm used [31,44].

However, GphT showed an antidepressant-like effect in the tail suspension test (TST) and forced swimming test (FST) since significantly reduced immobility time in both tests. The ability of haloperidol—a non-selective dopamine receptor antagonist— to reverse this effect suggests that GphT positively modulates the dopaminergic system. The doses of haloperidol used did not modify locomotor activity in the OFT, ruling out sedation as a possible bias in the TST. Pretreatment with ketanserin, a selective 5-HT2A receptor antagonist, prevented the antidepressant-like effect of GphT in the TST, suggesting a key involvement of 5-HT2A receptors in its serotonergic mechanism of action, consistent with previous reports for other plant-derived compounds [26, 27]. Similar to haloperidol, ketanserin did not alter locomotor activity in the open field test at doses used.

Since the influence of nitric oxide on antidepressant-like responses has been more consistently demonstrated in the forced swimming test (FST) than in the tail suspension test (TST), probably due to its greater sensitivity to serotonergic modulation [32], we evaluated the role of NO using the FST. Our results show that L-NAME and sildenafil, at subeffective doses reversed the antidepressant effect of GphT, suggesting the involvement of the nitric oxide (NO)/cyclic guanosine monophosphate (cGMP) signaling pathway. Sildenafil increases cGMP levels by inhibiting its degradation, which has been associated with depression-like behaviors [45]. Our result suggests that excess cGMP avoids the antidepressant mechanism of GphT. However, L-NAME, which also reversed the antidepressant effects of GphT, reduces the NO synthesis and therefore, it would be reducing the consequent activation of cGMP production. As a whole, these results suggest that a baseline level of NO is required for its antidepressant action, but an excess of the pathway NO/GMPc would become in the opposite effect. These findings indicate that GphT not only suppresses the NO/cGMP pathway but also modulates it within an optimal physiological range, since both excessive activation (by sildenafil) and excessive inhibition (by L-NAME) impair the antidepressant response. Therefore, the antidepressant mechanism of GphT likely depends on maintaining adequate NO levels and balanced NO/cGMP signalling.

Among the bioactive compounds identified in GphT, baicalin exhibits well-documented antidepressant-like, neuroprotective and anxiolytic-like effects in vitro and in vivo [39]. Moreover, apigenin glycosides, such as apigenin-7-O-glucoside, may contribute to antidepressant activity, as their aglycone, apigenin, has been shown to reduce depressive-like behavior in animal models through modulation of oxidative stress, inflammatory signaling, and cellular energy homeostasis in the hippocampus [46]. In contrast, gallic acid, (+)-catechin, p-coumaric acid, and acacetin derivatives identified in GphT have not been directly associated with antidepressant effects.

Taken together, these findings reveal that the neuropharmacological profile of Gomphrena perennis tincture is complex. While the extract contains compounds previously associated with anxiolytic, sedative, and antidepressant actions, in our experimental models GphT showed an anxiogenic response and antidepressant like-effect, without sedation. This pharmacological profile highlights the multifaceted interactions that can occur within complex phytochemical components, where their active components, their relative concentrations, and their potential synergistic or antagonistic effects determine the final pharmacodynamic outcome. In particular, nitric oxide (NO) levels appears to play a fundamental and paradoxical role in mediating these effects: inhibition of nitric oxide synthase by L-NAME prevented the anxiogenic response but also abolished the antidepressant effect, while potentiation of NO/cGMP signalling by sildenafil likewise prevented the antidepressant effect. These findings suggest that while maintaining nitric oxide in a regulated physiological range it promotes the antidepressant effect of GphT. However, these NO levels also contribute to the anxiogenic action of GphT, adding complexity to the potential therapeutic applications of this plant.

## 4. Materials and Methods

### 4.1. Preparation of the tincture of *G. perennis*

In spring, aerial parts of *G. perennis* L. were harvested from the Carlos Spegazzini Botanical Garden and Arboretum, School of Agricultural and Forestry Sciences of the National University of La Plata. The botanical identification was made plants by Marta Colares, MSc. The collected leaves were dried at 40 °C. The tincture of G. perennis (GphT) was prepared by macerating the dried leaves (20% w/v) in 70% v/v ethanol (yield = 7.1% w/w). For in vivo pharmacological experiments, GphT doses were expressed as mg of dried leaves/body weight of the animal (mg/kg).

### 4.2. Animals

In order to study the pharmacological activities of Gomphrena perennis tincture (GphT), male and female Swiss mice (25–30 g) were used. The animals were cared for according to international standards established by US guidelines (NIH publication #85-23, revised in 1996) and the principles in the Declaration of Helsinki. A local ethical committee of the School of Exact Sciences of the National University of La Plata approved the protocols (005-29-18; 002-43-19). Mice were only kept fasting for 24 hours for the novelty-suppressed feeding test (NSFT).

### 2.3. Acute toxicity study

The acute toxicity assessment of *Gomphrena perennis* L. tincture (GphT) was carried out in accordance with OECD Guideline 423 (Organization for Economic Cooperation and Development) [17]. For each dose level, three male mice were randomly selected. Prior to administration, the animals were fasted for at least four hours with free access to water. First, the GphT was administered intraperitoneally (i.p.) at a dose of 5 mg/kg, after which the animals were carefully observed for the first four hours to detect any sign of toxicity. Monitoring for potential mortality continued over the following three days. If two of the three mice died at this dose, the dose was classified as toxic. However, if only one died, the same dose was readministered to a new group of three mice. In the absence of mortality, higher doses (50, 300, and 2000 mg/kg) were subsequently tested to further evaluate the toxicity profile of GphT.

### 2.4. Behavioral evaluation in mice

All drugs were administered by intraperitoneal (i.p.) injections to mice in a volume of 0.1 ml per 30 g of body weight. The GphT doses used in the behavioural tests (400 and 800 mg/kg) were selected based on concentrations previously shown to be effective as antispasmodics in isolated rat intestine [6] and scaled according to estimated plasma levels. Since the dose of 2000 mg/kg did not show any sign of toxicity in the acute toxicity study, we considered the selected doses to be safe. Besides, the 400 mg/kg i.p. dose is consistent with doses used for a related species [18], while the dose of 800 mg/kg i.p. has been widely used to evaluate the central effects on other several plant species [19, 20].

The ethanolic vehicle (vehicle, assesed as negative control) was prepared with 70% v/v ethanol in the appropriate percentage as used for preparing the doses of 400 and 800 mg/kg of GphT respectively.

#### 2.4.1. Open field test (OFT)

Spontaneous locomotor activity, exploratory behavior, and emotional responses were assessed using the Open Field Test (OFT) as previously described. The apparatus consisted in a 30 × 50 x 27 cm white box, divided into 15 equal squares (10 cm² each) by black lines. Appropriate environmental conditions were maintained [21]. Before treatment, each animal was placed in the same designated corner of the field and allowed to explore freely for 5 minutes. The test was performed 60 minutes after receiving the GphT or vehicle. Diazepam (1 mg/kg), a known benzodiazepine, was used as the reference substance (positive control). The number of crossed lines, rearing, grooming behavior, and other relevant signs were recorded during each 5-minute session.

#### 2.4.2. Novelty-suppressed feeding test (NSFT)

Mice, housed in individual cages, were exposed to a 30-minute habituation period. One hour after intraperitoneal administration of the tincture or vehicle, each mouse was placed in the same corner of a wooden box (40 × 40 × 30 cm), which contained a white circular filter paper (11 cm in diameter) in the center of the box with a single food pellet. The latency to start eating was recorded. As soon as the animal took its first bite, it was transferred to its individual cage containing a pre-weighed food pellet. After 5 minutes, the amount of food consumed was measured. Then, mice were returned to their cages with free access to food and water [22]. Diazepam (0.3 mg/kg) was used as reference drug (positive control).

#### 2.4.3. Elevated plus-maze test (EPM)

To evaluate anxiety-like behaviors in Swiss mice, the Elevated Plus Maze (EPM) test was conducted as previously described [23]. The apparatus consists of a black wooden cross elevated to 100 cm above the floor, with four arms extending from a central platform. It consists of four arms extending from a central platform, two opposing open arms (30 × 5 cm each), without surrounding walls, while the other two are closed arms with walls arms (30 × 5 × 15 cm each) opposite to each other. For testing, mice were respectively treated via i.p. with vehicle, 0.3 mg/kg diazepam as anxyolitic reference substance, and GphT (400 and 800 mg/kg). An hour after the injection, the mice were placed on the central platform and the number of entries to the open and to the closed arms, and the time spent there, were counted during 5 min for each mouse.

#### 2.4.4. Tail suspension test (TST)

The tail suspension test (TST) was performed as previously described [24]. To assess the effects of GphT, a 50 cm high, 12 cm deep, and 60 cm wide setup was used. Mice were suspended individually in separate compartments to 10 cm from the floor, with adhesive tape placed 2 cm from the base of the tail. Measurements were made 1 hour after i.p. administration and videotaped for 6 minutes. Immobility was defined as the animal remaining in an upright position without hind limb movement. Immobility time (IT, in seconds) was recorded using a stopwatch. Clomipramine (1.25 mg/kg), a known tricyclic antidepressant, was used as the reference drug. To investigate the involvement of the monoaminergic system, mice were treated intraperitoneally (i.p.) with propranolol (1 mg/kg; a β-adrenergic receptor blocker; [25]), prazosin (1 mg/kg; an α1-adrenergic receptor antagonist [25, 26]), yohimbine (1 mg/kg; an α2-adrenergic receptor antagonist, [25, 26]), ketanserin (a 5-HT2A/2C receptor antagonist, [27]), ondansetron (a 5-HT3 receptor antagonist [28]), or haloperidol (a dopamine D1/D2 receptor antagonist, [27]), five min before vehicle or GphT (400 or 800 mg/kg, i.p.) administration; immobility time was measured 1 hour later. The antagonist doses used have been previously described as subeffective—they do not induce antidepressant-like effects on their own—and are employed as pharmacological tools to investigate the mechanisms of action involved. [25–28]. Moreover, in order to assess whether changes in locomotion could influence TST performance, other groups of mice similarly treated were subjected to a 6-minute OFT one hour after drug administration.

#### 2.4.5. Forced swimming test (FST)

The forced swimming test (FST) was performed according to previously established protocols. Swiss mice received intraperitoneal injections and, after 1 hour, were placed in a clear glass cylinder (24 cm in diameter and 22 cm in height) filled with water at a temperature of 22–25°C. Each session lasted 6 minutes, and immobility time (IT) was recorded during the last 4 minutes. Immobility was defined as passive floating of the animal, with only the minimal movements necessary to keep its head above water. After the test, the mice were dried and placed in a warm environment. The bath water was changed after every session to avoid influence on the next mouse [29]. Clomipramine (1.25 mg/kg) was used as the antidepressant reference drug. Similarly to the TST protocol, other group of mice were evaluated in the open field test for 6 minutes, one hour after intraperitoneal administration, to account for potential effects on locomotion that could influence FST results.

To investigate the involvement of the nitric oxide-cyclic guanosine monophosphate (NO/cGMP) pathway, mice were intraperitoneally (i.p.) treated with L-NAME (10 mg/kg; a non-selective nitric oxide synthase inhibitor: [30]) or sildenafil (5 mg/kg; a phosphodiesterase (PDE) inhibitor; [31]) 5 minutes before administration of vehicle or GphT (400 or 800 mg/kg, i.p.). Immobility time was measured one hour later. Similarly, another group of mice was treated with haloperidol prior to vehicle or GphT administration to evaluate the dopaminergic system in the FST. As for the TST, the antagonist doses used have been previously described as subeffective and are employed as pharmacological tools to investigate the mechanisms of action involved [30,31].

#### 2.4.6. Treatment and drugs

The drugs used in the biological tests were: diazepam (Roche, Argentina), clomipramine (Sandoz, Argentina), ketanserin tartarate (Sigma-Aldrich, USA), prazosin (Sigma-Aldrich, USA), yohimbine (Sigma-Aldrich, USA), haloperidol (Sigma-Aldrich, USA), propranolol (Sigma-Aldrich, USA), ondansetron (Zofran® 8 mg, solución inyectable); L-NAME (Sigma-Aldrich, USA), sildenafil (Saporiti, Argentina). Drugs were dissolved in saline and then administered by intraperitoneal route (i.p.) in a volume of 0.1 ml per 10 g of mouse body weight. The doses used of each drug were obtained from bibliographic data, as previously described in the sections above [25–31].

#### 2.4.7. Statistical analysis

Results were expressed as the mean ± SEM (standard deviation of the mean); n = number of experiments. Multiple comparisons were analyzed by one-way or two-way ANOVA of each groups of experiments as appropriate. Tukey’s paired post hoc tests were done between the multiple treatments, and their results are shown in each figure; P < 0.05 was considered significant. All statistical analyses were performed using GraphPad Prism 8.0 software.

## 5. Conclusions

In summary, this study provides the first preclinical evidence demonstrating that *Gomphrena perennis* tincture exerts effects on the central nervous system through a distinctive behavioral profile, characterized by anxiogenic and antidepressant activity, without sedative activity. While the coexistence of these effects may limit its immediate therapeutic potential, it also highlights the presence of bioactive compounds capable of modulating complex neurochemical pathways, including dopaminergic, serotonergic, and NO/cGMP signaling systems. These findings suggest that GphT acts as a multimodal neuromodulatory complex whose pharmacological outcome depends on the delicate balance between its components and their effects on interacting neurotransmitter systems.

The involvement of nitric oxide as a common mediator of signaling in both anxiogenic and antidepressant responses reinforces the hypothesis that GphT does not act through a single neurotransmitter system, but rather exerts a modulatory influence on interrelated pathways. Its ability to activate dopaminergic and serotonergic systems, while maintaining a delicate balance in NO/cGMP signaling, suggests a regulatory mechanism of central neurochemical processes.

## 6. Limitations of this study

This study has several limitations that should be addressed in future research. First, all experiments were performed using the whole *Gomphrena perennis* tincture (GphT), without evaluating isolated components. While whole-plant extracts are the most commonly used by people, this approach limits the ability to attribute the anxiogenic and antidepressant effects to specific bioactive molecules.

Therefore, future studies should aim to isolate and characterize the components responsible for the observed behavioral outcomes. From a translational perspective, the antidepressant properties highlighted the need to investigate isolated compounds and standardized fractions of GphT to elucidate the molecular determinants of this effect, reducing the anxiogenic component while maintaining its antidepressant effect. These efforts could contribute to the identification of phytochemical candidates relevant to the development of antidepressant drugs.

## Supplementary Materials

The following supporting information is available at Zenodo: https://doi.org/10.5281/zenodo.18033834. Appendix A1. Previously published chromatographic analysis of the *Gomphrena perennis* tincture used in this study. Figure A1: One-way ANOVA analysis of Figure 2 data; Table A1: One-way ANOVA analysis of Figure 2 data; Table A2: One-way ANOVA analysis of Figure 3 data; Table A3: Two-way ANOVA analysis of Figure 4 and Figure 5 data.

## Author Contributions

Conceptualization, M.I.R.; methodology, M.I.R. and A.E.C.; validation, A.M.B., G.G. and M.I.R.; formal analysis, A.M.B. and G.G.; investigation, A.M.B. and G.G.; resources, M.I.R.; data curation, A.M.B. and G.G.; writing—original draft preparation, M.I.R.; writing—review and editing, M.I.R. and A.E.C.; visualization, M.I.R.; supervision, M.I.R.; project administration, M.I.R.; funding acquisition, M.I.R.

A.M.B. and G.G. contributed equally to this work.

All authors have read and agreed to the published version of the manuscript.

## Funding

This research was funded by Universidad Nacional de La Plata, grant number PPID-X604-UNLP. The APC was funded by the authors.

## Data Availability Statement

The datasets generated and/or analyzed during the current study have been deposited in the RI CONICET Digital (CONICET Institutional Repository) under the title *“Efectos centrales de la tintura de Gomphrena perennis”.* Public access to the data will be enabled once the repository’s curation and validation process is completed.

## Acknowledgments

Figures of the graphic abstract created with BioRender.com

## Conflicts of Interest

The authors declare no conflicts of interest.

## Abbreviations

The following abbreviations are used in this manuscript:

GphT: Gomphrena perennis tincture
RP-HPLC-DAD: Reversed Phase High-Performance Liquid Chromatography with Diode Array Detection
NSFT: Novelty-suppressed feeding test
OFT: Open field test
EPM: Elevated plus-maze test
TST: Tail suspension test
FST: Forced swimming test

## Disclaimer/Publisher’s Note

The statements, opinions and data contained in all publications are solely those of the individual author(s) and contributor(s) and not of MDPI and/or the editor(s). MDPI and/or the editor(s) disclaim responsibility for any injury to people or property resulting from any ideas, methods, instructions or products referred to in the content.

## Notes

https://doi.org/10.5281/zenodo.18033834

### Competing Interest Statement

The authors have declared no competing interest.

